# Parallel spatial channels converge at a bottleneck in anterior word-selective cortex

**DOI:** 10.1101/508846

**Authors:** Alex L. White, John Palmer, Geoffrey M. Boynton, Jason D. Yeatman

## Abstract

In most environments, the visual system is confronted with many relevant objects simultaneously. That is especially true during reading. However, behavioral data demonstrate that a serial bottleneck prevents recognition of more than one word at a time. We used functional magnetic resonance imaging (fMRI) to investigate how parallel spatial channels of visual processing converge into a serial bottleneck for word recognition. Observers viewed pairs of words presented simultaneously. We found that retinotopic cortex processed the two words in parallel spatial channels, one in each contralateral hemisphere. Responses were higher for attended words than ignored words, but were not reduced when attention was divided, even though behavioral performance suffered greatly. We then analyzed two word-selective regions along the occipito-temporal sulcus (OTS) of both hemispheres (i.e., sub-regions of visual word form area, VWFA). Unlike retinotopic cortex, each word-selective region responded to words on both sides of fixation. Nonetheless, a single region in the left hemisphere (VWFA-1 in posterior OTS) contained spatial channels for both hemifields that were independently modulated by selective attention. Thus, the left posterior VWFA supports parallel processing of multiple words. In contrast, a more anterior word-selective region in the left hemisphere (VWFA-2 in mid-OTS) showed limited spatial and attentional selectivity, consistent with activity of a single channel. Therefore, the visual system can process two words in parallel up to a late stage in the ventral stream. The transition from two parallel channels to a single channel in more anterior regions is consistent with the observed bottleneck in behavior.

## Introduction

Pages of text are among the most complex and cluttered visual scenes that humans encounter. You cannot immediately comprehend the hundreds of meaningful symbols on this page because of fundamental limits to the brain’s processing capacity. How severe are those limits, and what causes them? Studies of eye movements during natural reading have fueled a long debate about whether readers process multiple words in parallel (Engbert, Nuthmann, Richter, & Kliegl, 2005; Kennedy, 2000; Murray, Fischer, & Tatler, 2013; Reichle, Liversedge, Pollatsek, & Rayner, 2009; Starr & Rayner, 2001). In a direct psychophysical test of the capacity for word recognition, we recently found that observers can report the semantic category of only one of two words that are briefly flashed and masked. Those data demonstrate that a serial bottleneck allows only one word to be fully processed at a time (White, Palmer, & Boynton, 2018). Where is that bottleneck in the brain’s reading circuitry?

Early stages of visual processing are spatially parallel. In retinotopic areas of the occipital lobe, receptive fields are small, such that neurons at different cortical locations simultaneously process objects at different visual field locations. The spatial selectivity of visual neurons allows spatial attention to prioritize some objects: activity is enhanced at the points on the cortex that represent task-relevant compared to irrelevant locations (Beck & Kastner, 2014; Brefczynski & DeYoe, 1999; Gandhi, Heeger, & Boynton, 1999). During simple feature detection tasks, multiple attended locations can be enhanced in parallel with no cost (White, Runeson, Palmer, Ernst, & Boynton, 2017).

It is not clear whether such parallel processing extends into the brain areas responsible for complex object recognition. Ventral occipito-temporal cortex (VOTC) contains a mosaic of regions that each respond selectively to stimuli of a particular category, such as faces, scenes, objects or words (Grill-Spector & Weiner, 2014). Receptive fields in VOTC span much of the visual field, so it is unclear how any one region can process multiple stimuli of its preferred category (Agam et al., 2010; Bao & Tsao, 2018; Gentile & Jansma, 2010). In the case of word recognition, nearly every neuroimaging study has presented only a single word at a time. Moreover, while there are detailed models of attention in retinotopic cortex, we know relatively little about the function of spatial attention in human VOTC (Kay, Weiner, & Grill-Spector, 2015; Reddy, Kanwisher, & VanRullen, 2009; Zumer, Scheeringa, Schoffelen, Norris, & Jensen, 2014). Of specific relevance to the present study, there have been no investigations of selective spatial attention in word-selective cortex. The limits of parallel processing and attentional selection have important implications for explaining limits on human behavior, especially during complex tasks such as reading.

We measured fMRI responses while observers performed a semantic categorization task (**Figure 1A**). On each trial, observers viewed a masked pair of words, on one each side of fixation, and either focused attention on one side (focal cue left or right) or divided attention between both sides (distributed cue). Of particular interest is a specialized VOTC region for word recognition, termed the “visual word form area” (VWFA) (Cohen et al., 2000; Dehaene, Le Clec, Poline, Le Bihan, & Cohen, 2002). Most authors refer to a single left-hemisphere VWFA that was originally proposed to be “invariant” to the visual field position of a word (Cohen et al., 2000, 2002; Dehaene et al., 2004; Price & Devlin, 2011). More recent fMRI studies, however, demonstrate that VWFA voxels have some spatial tuning, though are not organized retinotopically (Gomez, Natu, Jeska, Barnett, & Grill-Spector, 2018; Le, Witthoft, Ben-Shachar, & Wandell, 2017; Rauschecker, Bowen, Parvizi, & Wandell, 2012). Words also activate right hemisphere VOTC, and there may be a posterior to anterior hierarchy within word-selective cortex (Dehaene et al., 2004; Vinckier, Dehaene, Jobert, Dubus, & Sigman, 2007). Recent work has further identified two distinct sub-regions in the left hemisphere occipito-temporal sulcus (OTS) (Grill-Spector & Weiner, 2014; Stigliani, Weiner, & Grill-Spector, 2015). Compared to the more posterior region (VWFA-1), the anterior region (VWFA-2) is more sensitive to abstract lexical properties and is more strongly connected to language areas (Lerma-Usabiaga, Carreiras, & Paz-Alonso, 2018).

**Figure 1.**
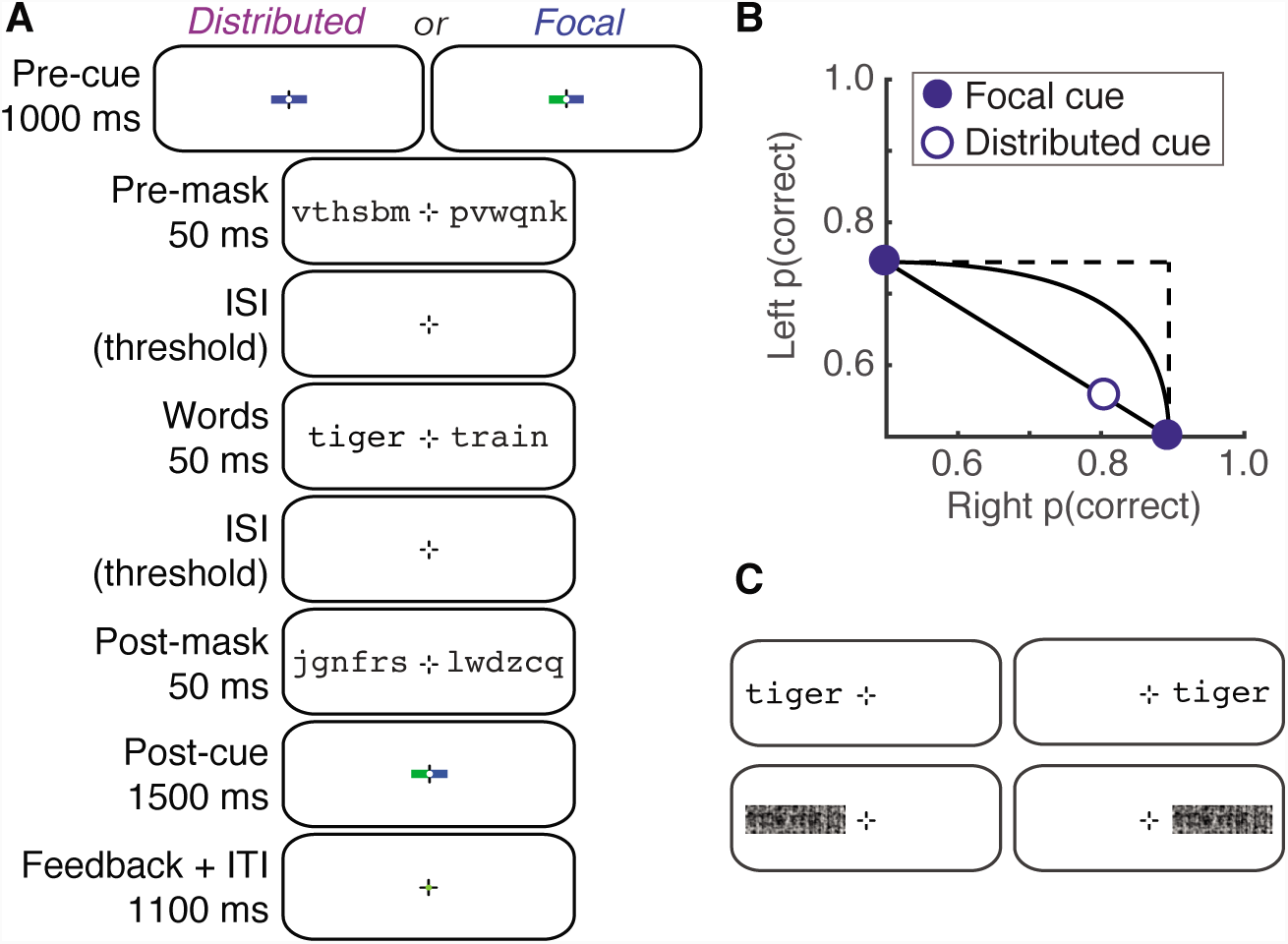
Stimuli and behavioral performance. **(A)** Example trial sequence in the main experiment. Half the participants attended to blue cues and half to green cues. Not shown are two additional blank periods with only the fixation mark: a 50 ms gap between the pre-cue and the pre-mask; and a gap between the post-mask and the post-cue, with duration set to 200 ms minus the sum of the two interstimulus intervals (ISIs) between the masks and words. **(B)** Mean semantic categorization accuracy collected in the scanner, plotted on an Attention Operating Characteristic. Error bars (±1 SEM, n=15) are so small that the data points obscure them. **(C)** Examples of the four stimulus conditions in the localizer scans (word-left, word-right, scramble-left, scramble-right).

With retinotopic visual cortex as a reference, we investigate whether the activity in each VWFA region is consistent with processing before or after the bottleneck that constrains word recognition. A region prior to the bottleneck should have two properties: (1) Its neurons (and voxels) should have sufficiently heterogeneous spatial tuning to process each word in a parallel spatial “channel”: a set of neurons that respond to stimuli at a particular visual field position. (2) Selective spatial attention should modulate the channels such that task-relevant words are prioritized prior to the bottleneck. In addition, a third property would support the hypothesis of *unlimited-capacity* parallel processing: little to no reduction in response when attention is divided between both words compared to focused on one. These three properties characterize the responses of retinotopic regions during a simple visual detection task (White et al., 2017). In contrast, a region in which BOLD responses reflect processing after the bottleneck must contain only a single channel. It therefore should have two properties: (1) more uniform spatial tuning; and (2) identical responses in each attention condition.

## Results

### Observers can recognize only one word at a time

In an fMRI experiment, observers viewed pairs of nouns, one on either side of fixation, which were preceded and followed by masks made of random consonants (**Figure 1A**). At the end of each trial, observers were prompted to report the semantic category (living vs. non-living) of one word. In the *focal cue condition,* a pre-cue directed their attention to the side (left or right) of the word they would need to report. In the *distributed cue condition,* a pre-cue directed them to divide attention between both words and at the end of the trial they could be asked about the word on either side. In a training phase, we set the duration of the inter-stimulus intervals (ISIs) between the words and the masks to each observer’s 80% correct threshold in the focal cue condition, and then maintained that timing for all conditions in the experiment. The mean (± standard error) ISI was 84±5 ms. We excluded trials with eye movements away from the fixation mark.

Accuracy was significantly worse in the distributed than focal cue condition: the mean difference in proportion correct was 0.14±0.01 (95% CI= [0.12 0.16], t(14)= 14.1, p<10 ^−7^). Accuracy was also significantly worse on the left than the right side of fixation, both in the focal cue condition (mean difference = 0.15 *±* 0.02; 95% CI=[0.12 0.19], t(14)=7.60, p<10^−7^) and in the distributed cue condition (0.25 ± 0.02; 95% CI= [0.20 0.29], t(14)=10.7, p<10^−5^). This hemifield asymmetry for word recognition is well established (Mishkin & Forgays, 1952; White et al., 2018).

In **Figure 1B** we plot these data on an “Attention Operating Characteristic” (AOC; Sperling & Melchner, 1978). Accuracy for the left word is plotted against accuracy for the right word. The focal cue conditions are pinned to their respective axes. The distributed cue condition is represented by the open symbol. Also shown are the predictions of three models for where that point should fall (Bonnel & Prinzmetal, 1998; Scharff, Palmer, & Moore, 2011; Shaw, 1980; Sperling & Melchner, 1978):

1. *Unlimited-capacity parallel processing:* two words can be fully processed simultaneously just as well as one. This predicts that the distributed-cue point falls at the intersection of the dashed lines (no deficit), as has been found for simpler visual detection tasks (Bonnel, Stein, & Bertucci, 1992; Scharff et al., 2011; White et al., 2018, 2017)
2. *Fixed-capacity parallel processing:* The brain extracts a fixed amount of information from the whole display per unit time, using processing resources that must be shared between both words. Varying the proportion of resources given to the right word traces out the black curve in the AOC (White et al., 2018).
3. *All-or-none serial processing:* words are recognized one at a time, and because of the time constraints imposed by the masking, only one word can be processed per trial. Varying the proportion of trials in which the right word is processed traces out the diagonal black line in the AOC.

Mean dual-task accuracy fell significantly below the predictions of both the unlimited-and fixed-capacity parallel models, and perfectly on top of the all-or-none serial model’s prediction (**Figure 1B**). The average minimum distance from the serial model’s line (calculated such that points below the line have negative distances) was 0.0±0.01 (95% CI=[−0.027 0.027]; t(14)=0.10, p=0.925). In sum, when observers tried to divide attention between the two words, they were able to accurately categorize one word (∼ 80% correct), but were at chance for the other. This behavior is consistent with the presence of a serial bottleneck at some point in the word recognition system (White et al., 2018).

### Selectivity for the contralateral hemifield decreases from posterior to anterior regions

We analyzed BOLD responses in retinotopic visual areas (V1-hV4, VO, LO), and several word-selective regions (VWFA-1 and VWFA-2; **Figure 2A**). In the left hemisphere, all participants had VWFA-1 in the posterior OTS as well as the more anterior VWFA-2. 14 of 15 participants also had a right hemisphere region symmetric to VWFA-1 but, only 5/15 participants had a more anterior right VWFA-2, consistent with prior work (Grill-Spector & Weiner, 2014; Strother, Coros, & Vilis, 2016).

**Figure 2.**
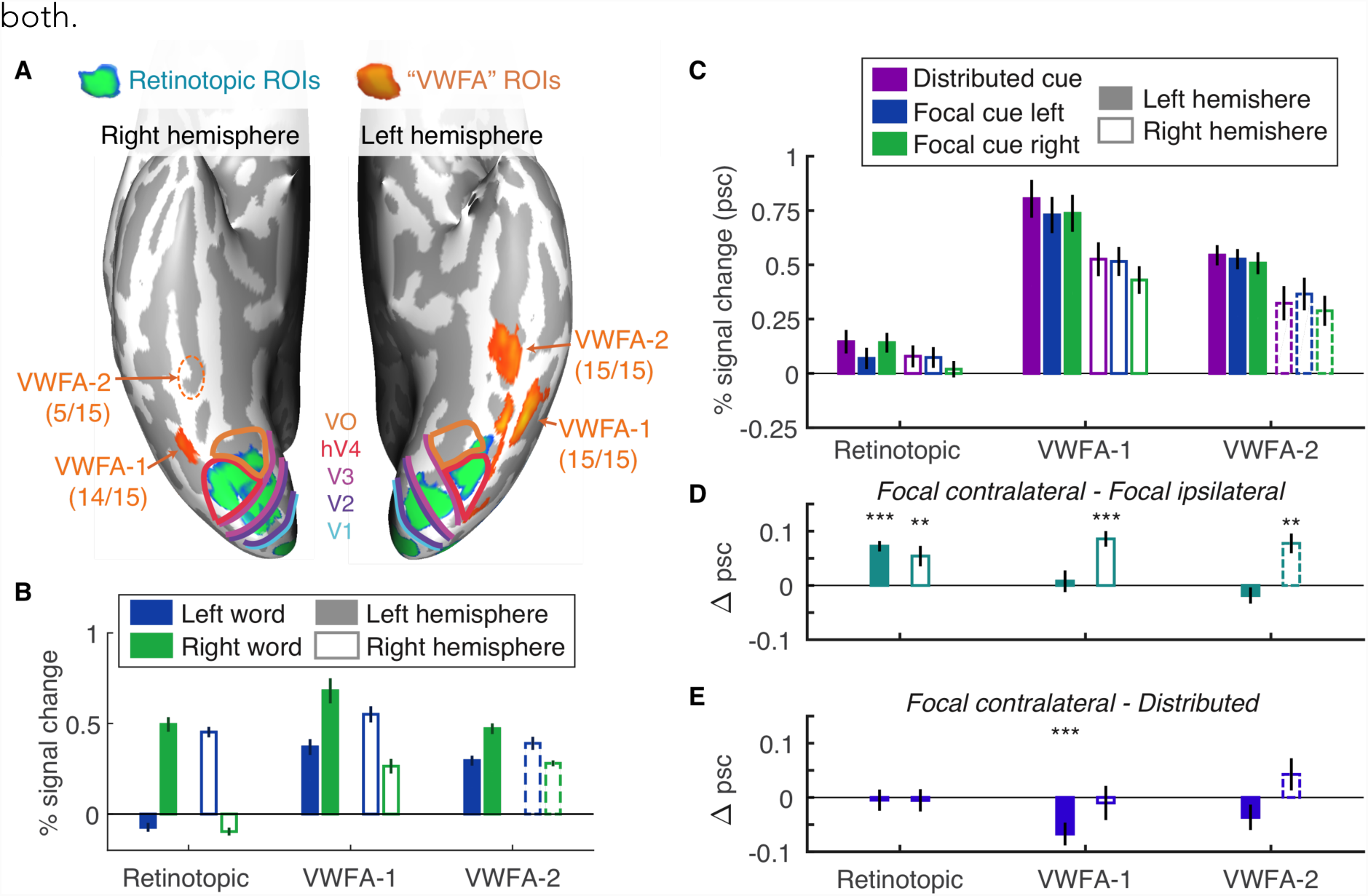
ROIs and mean BOLD responses. **(A)** Ventral view of the inflated cortical surfaces of one representative subject’s brain. Colored lines are the boundaries between retinotopic areas. The parenthetical numbers below each VWFA label indicates the number of subjects in which that area could be defined. Consistent with previous studies, we find right VWFA-2 in a minority of subjects, and represent its data with dashed lines to indicating that it is not representative of the average subject. **(B)** Mean responses to words on the left and right of fixation during the localizer scans. **(C)** Mean BOLD responses in the main experiment, divided by region, hemisphere, and pre-cue condition. **(D)** Mean selective attention effects: differences between responses when the contralateral vs. ipsilateral word was focally cued. **(E)** Mean divided attention effects: differences between responses when the contralateral word was focally cued vs. both words were cued. Error bars are ± 1 SEM. Asterisks indicate two-tailed p-values computed from bootstrapping: *** = p<0.001; ** = p<0.01; * = p<0.05.

In order to assess how each region processes pairs of words that are presented simultaneously, we first need to know the region’s sensitivity to single words at different locations in the visual field. We analyzed the mean responses to single words presented at either the left or right location in the localizer scan (stimuli in **Figure 1C**, results in **Figure 2B**). Consistent with the well-known organization of retinotopic cortex, the left hemisphere retinotopic areas responded positively only to words on the right of fixation, and vice versa.

The VWFAs, in contrast, are only partially selective for the contralateral hemifield, and that hemifield selectivity decreases from VWFA-1 to VWFA-2 (**Figure 2B**). Although all the VWFAs respond positively to words on both sides of fixation, most voxels still prefer the contralateral side (**Supporting Information Appendix Figure S1**). We assessed lateralization with the index: LI = 1-R_i_/R_c_, where R_i_ and R_c_ are the across-voxel mean responses to ipsiiaterai and contralateral words, respectively (Rauschecker et al., 2012). Across subjects, the mean LI values were: 0.46±0.04 in left VWFA-1, 0.36±0.06 in left VWFA-2; 0.52±0.08 in right VWFA-1; and 0.27±0.03 in right VWFA-2. LI differed significantly between VWFA-1 and VWFA-2 (F(1,45)=6 71, p=0.012). That difference did not interact with hemisphere (F(1,45)=0.92, p=0.34), and was present when the left hemisphere was analyzed separately (F(1,28)=5.33, p=0.029). Note that the comparison of responses to words at the left and right locations was independent of the (words – scrambled words) contrast used to select the VWFA voxels (see **Methods**).

In summary, retinotopic areas selectively process words in the contralateral visual field. In contrast, the VWFAs of both hemispheres respond to single words at both locations, but with a preference for the contralateral side. That preference is weaker in VWFA-2 than VWFA-1, suggesting more integration across visual space. The magnitude of contralateral preference for single words, however, does not indicate whether either area could process two words at once in the main experiment. For example, right hemisphere VWFA-1 might process the left word while left VWFA-1 processes the right word (similar to right and left V1), or either area could process both words in parallel. We investigated those questions by analyzing data from main experiment, when two words were present simultaneously and the observer attended to one or both.

### Left VVVFAs respond strongly during the semantic categorization task

**Figure 2C** plots the mean BOLD responses in each ROI and cue condition, averaging over all the retinotopic areas (restricted to the portions that are sensitive to the locations of the words). See **Figure S2** for each retinotopic ROI separately. The VWFAs responded more strongly to the briefly flashed words than retinotopic regions, and the left hemisphere responded more strongly than the right, especially in the VWFAs. A linear mixed-effects model (LME) found reliable effects of region (F(7,639) = 156.9, p< 10 ^−133^), hemisphere (F(1,639) = 52.3, p< 10 ^−11^), and cue (F(2,639) = 9.2, p=0.0001). The effect of hemisphere interacted with region (F(7,639) = 3.98, p=0.0003), but no other interactions were significant (Fs<0.25).

### Hemispheric selective attention effects are reliable in retinotopic cortex but not in the left VWFAs

The behavioral data demonstrate that subjects cannot recognize both words simultaneously. In the focal cue condition, therefore, the mechanisms of attention must select the relevant word to be processed fully. We first assess the selective attention effect in each region by comparing the mean BOLD responses when the contralateral vs. ipsilateral side was focally cued (**Figure 2D**). No prior study has investigated such effects in the VWFAs.

An LME model found a main effect of cue (contralateral vs. ipsilateral; F(1,426) = 12.58, p=0.0004) that did not interact with region or hemisphere (Fs<0.5). We also conducted planned comparisons of the focal cue contralateral vs. ipsilateral responses in each ROI (**Figure 2D**). The selective attention effect was reliable in the retinotopic ROIs (mean effect: 0.06±0.01% signal change) and in right hemisphere VWFA-1 (0.09±0.01%) and VWFA-2 (0.08±0.02%). However, it was absent in left hemisphere VWFA-1 (0.01 ±0.02%) and VWFA-2 (−0.02±0.02%). We propose two explanations for the lack of effects in the left VWFAs: 1. those areas process both words, so the mean response is a mixture of attended and ignored words; or 2. they process only one attended word, which makes the mean BOLD response identical in the focal cue left and right conditions. The spatial encoding model described below allows us to discriminate between those possibilities.

### Mean BOLD responses show no evidence of capacity limits

Assuming that the BOLD response to each word is proportional to the signal-to-noise ratio of the stimulus representation, a region with a capacity limit should respond less strongly when attention is divided than focused. **Figure 2E** plots the mean divided attention effects, which are the differences between the focal cue contralateral and the distributed cue conditions. There was no main effect of cue or interaction with region or hemisphere (all Fs<0.5). Bootstrapping on each ROI found no significant divided attention effect except for an *inverse* effect (distributed > focal) in left VWFA-1 (mean: 0.07 ± 0.02% signal change). Therefore, this analysis revealed no evidence of a capacity limit in any area. The data are similar to those in a previous study that found showing unlimited capacity processing of simple visual features in retinotopic cortex (White et al., 2017).

### Left VWFA-1 contains two parallel channels that are modulated by selective spatial attention

Because the VWFAs respond to words on both sides of fixation, we cannot isolate the response to each word within them simply by computing the mean response in the contralateral region (as we can do for retinotopic regions). Instead, we capitalize on differences in spatial tuning across individual voxels and build a “forward encoding model” (Brouwer & Heeger, 2009; Sprague et al., 2018). The model assumes that there are (at least) two spatial “channels” distributed across the region: one for the left word and one for the right word. Each voxel’s response is modeled as a weighted sum of the two channel responses. We estimate the two weights for each voxel as its mean responses to single words on the left and right in the localizer scans. We then “invert” the model via linear regression to estimate the two *channel responses* in each condition of the main experiment (**Figure 3A**). Comparing channel responses across cue conditions indexes the effect of spatial attention on voxels that are tuned to specific locations and reveals effects that were obscured by averaging over all voxels in an ROI.

**Figure 3.**
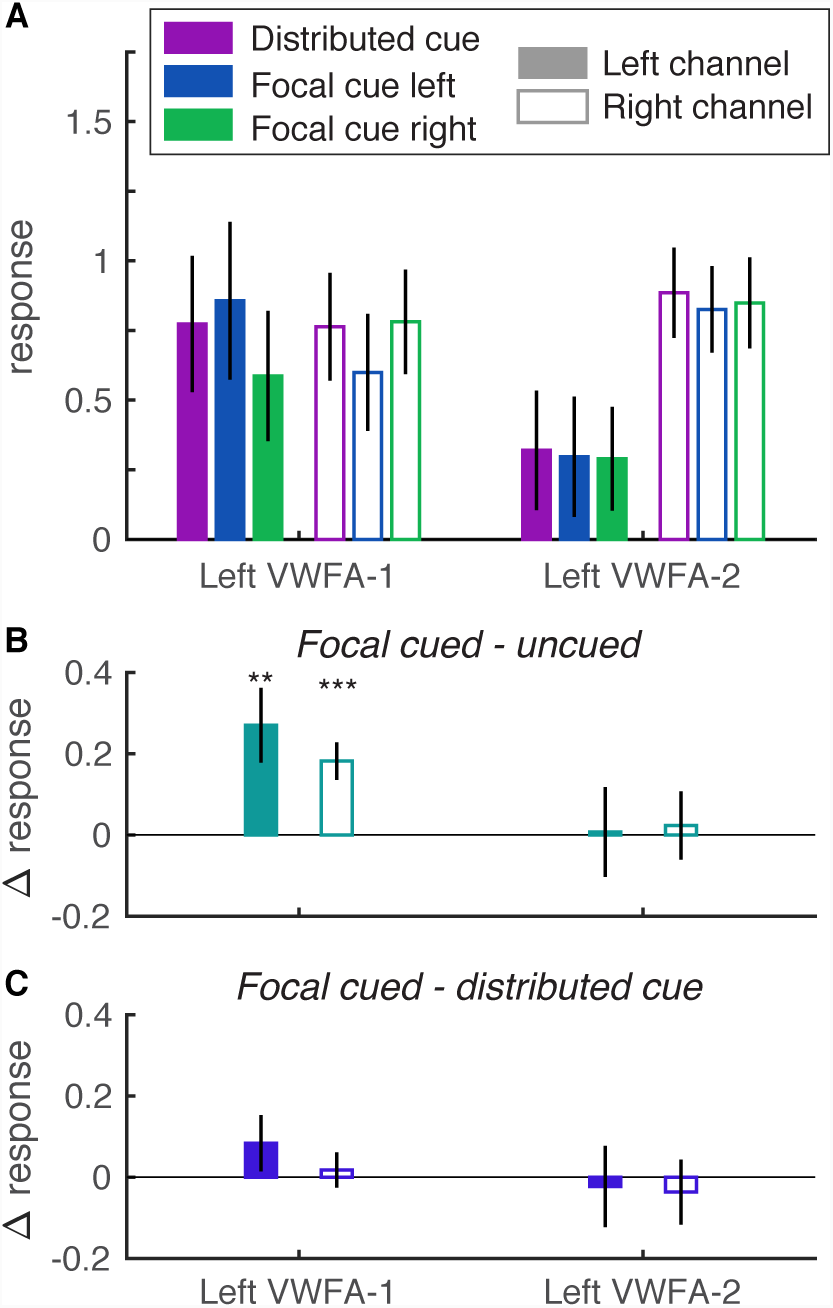
Estimated left hemisphere channel responses from the spatial encoding model. (**A**) Mean responses, separately for each ROI, channel, and cue condition. The left channel is plotted with solid bars and the right channel with open bars, while the bar colors indicate pre-cue conditions. (**B**) Selective attention effects: the differences between each channel’s responses when its visual field location was focally cued vs. uncued. (**C**) Divided attention effects: the differences between each channel’s response when its location was focally cued vs. when both sides were cued. Error bars and asterisks as in Figure 2. See Figure **S4** for right hemisphere data.

But not all regions necessarily contaIn two parallel spatial channels; in fact, we predict that a region after the bottleneck should only have one channel. Therefore, we also fit a simpler one-channel model to each region and compared its fit quality to the two-channel model. In the one-channel model, each voxel is given a single weight: the average of its localizer responses to left and right words. We then model the voxel responses In each condition of the main experiment by scaling those weights by a single channel response.

Adjusting for the number of free parameters, we found that the two-channel model fit significantly better than the one-channel model in left hemisphere VWFA-1: mean adjusted R^2^ = 0.63 vs. 0.57; 95% Cl of difference between models = [0.02 0.19], In left VWFA-2, the two-channel model fit slightly worse than the one-channel model: 0.40 vs. 0.36; 95% CI of difference = [−0.14 0.020). In right hemisphere VWFA-1 and VWFA-2, the one-channel model fit significantly better (see **Figure S3**). We therefore only reject the one-channel model for left hemisphere VWFA-1. Given that not all subjects have right hemisphere VWFAs, and the one-channel model fit significantly better there, we plot the estimated responses from the two-channel model only for the left hemisphere in **Figure 3** For the right hemisphere, see **Figure S4**

We measured selective attention effects within each ROI as the difference between each channel’s responses when its preferred location was focally cued vs. uncued (**Figure 3B**). We fit LMEs to assess those cue effects and how responses differed across the left and right channels. In Left VWFA-1, there was a main effect of cue (mean: 0.23 ± 0.07; F(1,56)= 12.3, p=0.001), no main effect of channel (F(1,56)=0.01, p=0.94), and no interaction (F(1,56)=2.41, p=0.13). The selective attention effect was significant in both channels. The average cued response was 1.38 times the average uncued response. Left VWFA-2 showed a very different pattern, with no significant effects of cue (mean: 0.02±0.09; F(1,56)=0.03, p=0.87), or channel (mean right-left difference: 0.55±0.34; F(1,56)=2.88, p=0.10), and no interaction (F(1,56)=0.09, p=0.77). This pattern is consistent with the observation that the one-channel model is adequate for VWFA-2.

Channel responses in the right hemisphere VWFAs (**Fig. S4**) partially matched what was observed in the left hemisphere. Only the left channel within right VWFA-1 had a significant spatial attention effect. More detail is provided in the SI, but note that the one-channel model was the best fit for right hemisphere areas, so there is limited value in interpreting those data. In summary, only in left VWFA-1 did we find evidence of two parallel channels, within a single brain region, that could both be independently modulated by selective spatial attention.

### Spatial and attentional selectivity are correlated in VWFA-1

We performed one more test of whether each region supports parallel spatial processing and attentional selection prior to the bottleneck. If so, the magnitude of the spatial attention effect in each voxel should be related to its spatial selectivity. Consider a voxel that responds equally to *single* words on the left and right. When two words are presented at once, attending left would affect the voxel response in the same way as attending right. This voxel with no spatial selectivity should therefore have no selective attention effect. In contrast, a voxel that strongly prefers single words on the right should respond much more in the focal cue right than focal left condition. We tested this prediction by evaluating the linear correlation between two independent measures from individual voxels in separate scans: (a) the difference between responses to single words on the left and right; (b) the difference between responses in the focal cue left and right conditions. That correlation was significantly positive for all ROIs except left VWFA-2 (**Figure S5**). This means that VWFA-1 behaves like other visual areas: the differential spatial tuning of its voxels (and neurons) allows parallel processing of items at different spatial locations, and attentional selection of task-relevant items. This also applies to the right hemisphere VWFAs, which primarily process the left visual field. The one-channel model nonetheless fit them best because they do not simultaneously represent both the left word and the right word. Only left VWFA-1 appears capable of doing that.

Finally, left VWFA-2 is unique in that attention effects on individual voxels (which average to 0) are not related to their spatial preferences. This result further supports the hypothesis that left VWFA-2 represents a single word after the bottleneck, perhaps in a more abstract format that can be communicated to language regions.

### Differences between VWFA-1 and VWFA-2 do not reflect SNR

The differences between left VWFA-1 and VWFA-2 could be due to lower signal-to-noise of our measurements in VWFA-2. Specifically, it is plausible that increased noise obscured the presence of two channels and attention effects in VWFA-2. To test this possibility, we examined the patterns of spatial selectivity in the localizer scan data. Under the one-channel model, voxels may respond more to words in the contralateral visual field on average (**Fig. 1B**), but *differences* between the spatial preferences of individual voxels are just noise. In contrast, the two-channel model assumes that voxels within a region differ meaningfully in how much they prefer the left or right side (because the two words are processed in partially separable populations of neurons).

The two models make different predictions for the across-voxel correlation between responses to single words on the left *(W*_*L*_) and on the right *(W*_*R*_). All else being equal, the correlation should be *weaker* in a region that contains two channels than a region with only one channel, because of the variance added by the true differences in voxel preferences. The correlation should also be weaker in an area with more measurement noise in the BOLD response. So, if VWFA-2 has two channels that are obscured by additional measurement noise, its mean correlation coefficient between *W*_*L*_ and W_R_ must be *lower* than in VWFA-1. However, the opposite was true: mean r=0.72±0.09 in left VWFA-1 and r=0.81±0.04 in left VWFA-2. Moreover, as predicted by the additional heterogeneity in voxel preferences in a two-channel area, the average standard error of differences between *W*_*L*_ and W_R_ was greater in left VWFA-1 (0.27±0.04) than VWFA-2 (0.13±0.01). Therefore, measurement noise alone cannot account for the different pattern of results in VWFA-2. Our interpretation is that two spatial channels in VWFA-1 merge into a single channel in VWFA-2.

### Divided attention does not significantly reduce VWFA channel responses

Finally, we assessed the divided attention effects in each region by comparing the responses of each channel when its location was focally cued vs. when both locations were cued. The divided attention effects for the left hemisphere are plotted in **Figure 3C**. There was no effect of cue or interaction between cue and channel (all ps>0.20). For data from the right hemisphere (where the one-channel model was the best fit), see **Figure S4C**.

### Neuronal Attention Operating Characteristics (AOCs) assess capacity limits

We introduce a new analysis of BOLD data: a neuronal AOC (**Figure 4**) that can assess capacity limits similarly to the behavioral AOC (**Figure 1 B**). For a related analysis of EEG data, see (Mangun & Hillyard, 1990). In the behavioral AOC, the points pinned to the axes are the focal cue accuracy levels, relative to the origin of 0.5, which is what accuracy would be for an ignored stimulus. Correspondingly, on the neuronal AOC, we plot the differences between responses to attended and ignored words, with the right word on the x-axis and the left word on the y-axis. The solid points on the axes are the differences in response to each stimulus when it was focally cued vs. uncued. The single open point represents the distributed cue condition: its x-value is the difference in right word response between the distributed cue and focal *uncued* (i.e., focal cue left) conditions. Similarly, the y-value is the difference in left word response between the distributed cue and focal uncued conditions. We compare that distributed-cue point to the predictions of two models:

1. *Unlimited capacity parallel processing: as* in the behavioral AOC, this model predicts that the distributed-cue point falls on the intersection of the two dashed lines. That indicates no change in response magnitudes relative to when each word was focally cued.
2. *Serial switching of attention:* the behavioral data suggest that on each distributed-cue trial, observers recognize one word but not the other. It is as if they pick one side to attend to fully and switch sides sporadically from trial to trial. This model predicts that their brain state should be a linear mixture of the focal-cue left state and the focal-cue right state. That prediction corresponds to the diagonal line connecting the two focal-cue points.

**Figure 4** contains the averaged AOCs constructed from data averaged over all 15 observers. For retinotopic cortex, the mean data were clearly consistent with the unlimited-capacity model, as the distributed cue point fell just above the dashed intersection (**Fig. 4A**). We also assessed the distribution of AOC points across individual observers. The individual data in retinotopic cortex were limited by noise, as these areas were hardly responding above baseline (**Fig. 2C**). We could not construct the AOC for 4 of the 15 observers because they lacked a positive selective attention effect in at least one hemisphere, which put the whole AOC below one axis and rendered it uninterpretable. Among the remaining 11 observers, the mean distance from the nearest point on the serial switching line (calculated such that points below the line are negative) was: 0.05 ± 0.03 (95% CI = [−0.02 0.09]). We also computed the distance of each observer’s distributed-cue point from the unlimited-capacity parallel point: mean = 0.0 ±0.03 (95% CI = [−0.07 0.04]). That distance was negative if, averaged across hemispheres, distributed cue responses were less than focal cued responses. In sum, we cannot definitively rule out the serial-switching model for retinotopic cortex, but average responses were more consistent with the unlimited-capacity parallel model (**Fig 4A**).

**Figure 4.**
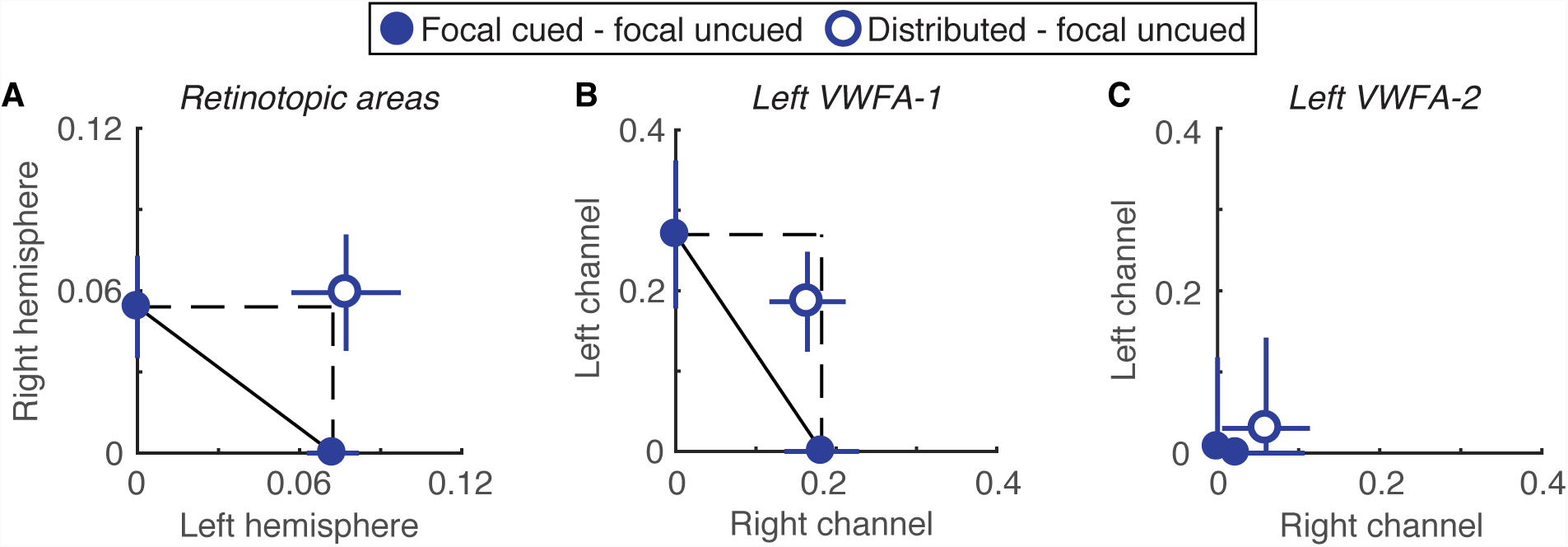
Neuronal Attention Operating Characteristics. **(A)** BOLD responses averaged across all retinotopic ROIs and all observers. The left hemisphere, which processes the right word, is plotted on the horizontal axis, and the right hemisphere is on the vertical axis. **(B)** Channel responses in left hemisphere VWFA-1. The channel for the right word is on the horizontal axis and the left channel is on the vertical axis. **(C)** Channel responses in left VWFA-2. Right hemisphere VWFAs are not plotted because the 1-channel model fit their responses better, and a minority of subjects had a right VWFA-2.

The AOC for channel responses in left VWFA-1, averaged over all 15 observers, is shown in **Figure 4B**. We were also able to construct these AOCs for 13 individuals. The mean distance from the nearest point on the serial switching line was 0.08 ± 0.04. The 95% confidence interval on that distance excluded 0: [0.02 0.16]. The mean distance of the distributed cue point from the unlimited capacity point was −0.11 ± 0.09, and not significantly different from 0 (95% confidence interval: [−0.26 0.06]). In sum, although there was a modest reduction channel responses when attention was divided, we can reject the serial switching model for left VWFA-1. That result suggests that the computations carried out in left VWFA-1 occur prior to the serial bottleneck that constrains recognition accuracy.

In left VWFA-2, the AOC collapses to the origin **(Figure 4C)** because responses were approximately equal under all cue conditions. This supports the hypothesis that left VWFA-2 responds to just one attended word, regardless of location. Indeed, two spatial channels are not necessary to explain the voxel responses in that region.

## Discussion

### A bottleneck in the word recognition circuit

The primary goal of this study was to determine how the neural architecture of the visual word recognition system forms a bottleneck that prevents observers from recognizing two words at once. Activity in retinotopic cortex matched three criteria for parallel processing prior to the bottleneck: (1) the two words were processed in parallel spatial channels, one in each cerebral hemisphere; (2) attended words produced larger responses than ignored words; (3) responses were equivalent when attention was divided between two words and focused on one word. These data support unlimited-capacity processing and are summarized in the neuronal Attention Operating Characteristic **(Figure 4A)**. We found a similar pattern in a prior study of retinotopic cortex with a simpler, non-linguistic visual task in which accuracy was the same in the focal and divided attention conditions (White et al., 2017). The fact that there was a severe (completely serial) divided attention cost to accuracy in this semantic categorization task demonstrates that attentional effects in retinotopic cortex do not always predict behavior.

Critically, a word-selective region in the left posterior OTS (VWFA-1) also supported parallel processing prior to the bottleneck. This single region is not retinotopically organized and responds to words on both sides of fixation. Nonetheless, its individual voxels are spatially tuned to different locations in the visual field (Le et al., 2017; Rauschecker et al., 2012). Here we demonstrate the functional significance of that tuning: we were able to recover the responses to both simultaneously presented words in parallel spatial channels within left VWFA-1. Those channels were independently modulated by spatial attention, and as shown in the AOC **(Figure 4B)**, the modest reduction caused by dividing attention was not sufficient to prevent parallel processing.

Finally, a relatively anterior word-selective region in the left hemisphere (VWFA-2 in the mid-OTS) had properties consistent with serial processing of single words after the bottleneck. It had weaker and more homogenous spatial selectivity, responded identically in all conditions of spatial attention, and its responses could be explained by a model with only one channel. Thus, compared to adjacent retinotopic areas and to VWFA-2, left hemisphere VWFA-1 is unique in having intermingled spatial channels covering both visual hemifields. Despite their spatial proximity and similar category selectivity, VWFA-1 and VWFA-2 therefore play distinct roles in the visual word recognition system.

On the basis of these findings, we propose the following model for how information flows through the word recognition circuit **(Figure 5)**. Visual signals from the retinas are first projected to contralateral retinotopic areas. Information about the left hemifield in right cortex then crosses over to left VWFA-1, presumably through the posterior corpus collosum. In the transition between VWFA-1 and VWFA-2 (or perhaps within VWFA-2 itself), there is a bottleneck, such that only one word can subsequently reach to higher-level language and decision areas. Spatial attention can boost one relevant word prior to the bottleneck to increase the likelihood that it is fully processed.

**Figure 5.**
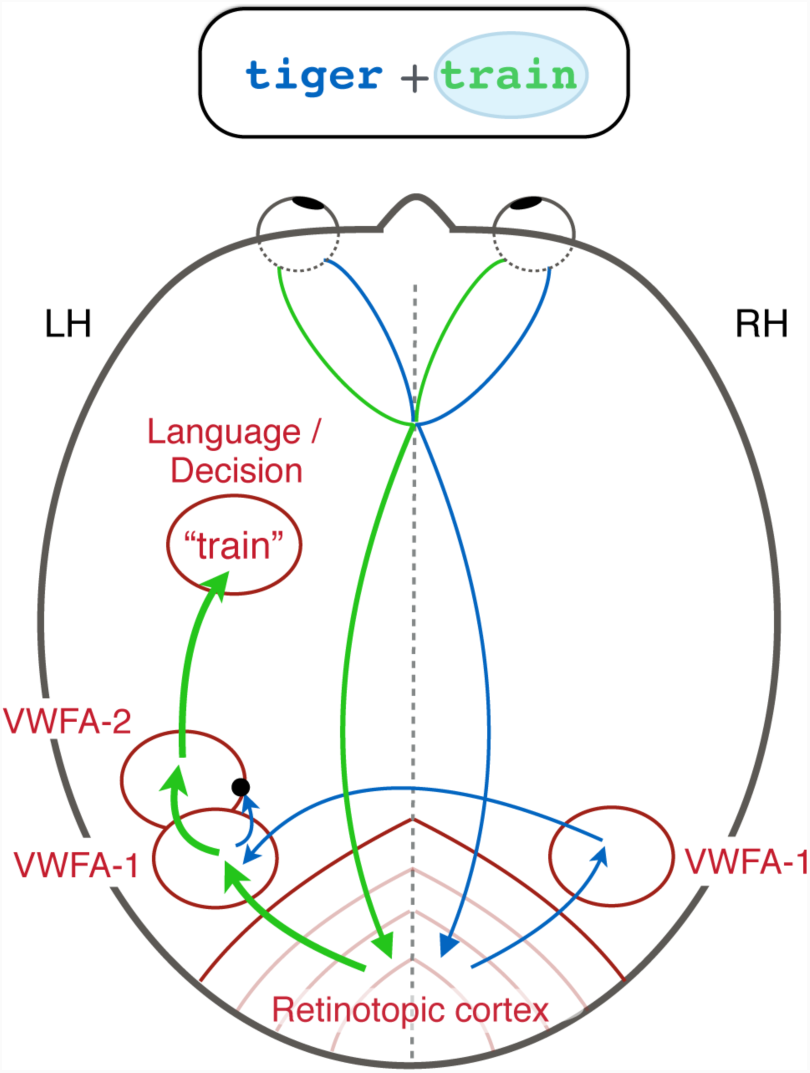
Circuit diagram of visual processing of two words. This brain is viewed from above, so the left hemisphere is on the left. The bubble around the word on the right side of the display indicates that it is selectively attended, and therefore its representation is relatively enhanced (thicker arrows). A bottleneck (black dot) prevents the unattended word from getting into left VWFA-2.

### Hierarchical processing and white matter connectivity

The differences we found between VWFA-1 and VWFA-2 build on previous models of the visual word recognition system. Several studies have concluded that the anterior portion of word-selective VOTC is more sensitive to higher-level, abstract, lexical properties (Dehaene et al., 2004; Lerma-Usabiaga et al., 2018; Vinckier et al., 2007). Another research group studied how VOTC integrates both halves of a single word that are split between hemifields (Strother et al., 2016; Strother, Zhou, Coros, & Vilis, 2017). They found that left VWFA-1 (which they label the “occipital word form area”) represented both halves of a word but maintained them separately. In contrast, a more anterior area (presumably VWFA-2) responded to entire words more holistically. That is consistent with our conclusion that left VWFA-1 contains two spatial channels while VWFA-2 contains only a single channel. Strother and colleagues also found that the right VWFA-1 was biased for the left hemifield, which is consistent with our finding that right VWFA-1 contained a single channel and was especially responsive when attention was focused to the left. Our results go further to show how this circuit responds to pairs of whole words, and to relate the multi-voxel patterns to selective attention and task performance.

Finally, our findings neatly align with recently discovered differences between VWFA-1 and VWFA-2 in terms of tissue properties and white matter connections (Lerma-Usabiaga et al., 2018). The more posterior OTS region is strongly connected through the vertical occipital fasciculus to the intraparietal sulcus, which is implicated in attentional modulations (Kay & Yeatman, 2017). That could explain why spatial attention modulates VWFA-1 but not VWFA-2. The mid-OTS region is connected through the arcuate fasciculus (Weiner, Yeatman, & Wandell, 2017) to temporal and frontal language regions. Lerma-Usabiaga et al. postulated that VWFA-2 “is where the integration between the output from the visual system and the language network takes place” (Lerma-Usabiaga et al., 2018). According to the present results, that output has capacity for only one word.

### Limitations and further questions

There are some limits to our interpretations. First, we were limited by the spatial resolution of our imaging technique, with 3×3×3 mm functional voxels. We found little evidence for multiple spatially tuned channels in the left VWFA-2, but it is possible that there are indeed sub-populations of neurons with different spatial tuning that are more evenly intermingled within voxels than in VWFA-1. Second, it is possible that two words are in fact represented separately in left VWFA-2, but in channels that are not spatially tuned. It is difficult to imagine how such an architecture would avoid interference between the two words, given that the observer must judge them independently and location is all that differentiates them.

Regarding left VWFA-1, the two-channel spatial encoding model may seem to imply that some neurons in that region are totally selective for the left visual field location, and others for the right. That is uncertain; indeed, 92% of *voxels* in left VWFA-1 responded more strongly to the right location than the left **(Figure S1)**. That is consistent with recent population receptive field measurements in VOTC (Le et al., 2017). But given that the 2-channel model fit best, we suppose that many voxels contain some neurons with receptive fields shifted far enough to the left that, when the subject attends to the left, they are up-regulated so left word is represented most strongly. The correlation between spatial preference and selective attention effects supports this interpretation **(Figure S5)**.

The conclusion that retinotopic areas have no capacity limit rests on an assumption that is common in the literature but deserves further scrutiny. Specifically, it assumes that the magnitude of the BOLD response is proportional to the signal-to-noise ratio of the stimulus representation used to make the judgment (Boynton, Demb, Glover, & Heeger, 1999; Ress, Backus, & Heeger, 2000). Our data showed that responses in the distributed cue condition were not reduced compared to focally cued stimuli. However, the BOLD signal may be affected by factors other than spatial attention that differ between cue conditions. For instance, the distributed condition was more difficult, so observers may have been more aroused. Some studies suggest that the total BOLD signal is a mixture of factors related to the stimulus, the percept, attention, anticipation, arousal, and perhaps other factors time-locked to the task (Cardoso, Sirotin, Lima, Glushenkova, & Das, 2012; Jack, Shulman, Snyder, McAvoy, & Corbetta, 2006; Ress & Heeger, 2003; Sirotin, Cardoso, Lima, & Das, 2012). We are aware of no prior results that could specifically explain the lack of divided attention effects in our data. But in principle, the total BOLD response could have been elevated by factors related to task difficulty while the strength of the stimulus representation was actually lowered in the distributed cue condition. Note, however, that no such factors could explain the *selective* attention effects, which are measured in trials with only focal cues. Our core conclusions about the parallel channels in VWFA-1 that converge in VWFA-2 hold even if we exclude the distributed-cue condition.

A final caveat is that the task we used differs markedly from natural reading. Specifically, observers fixated between two unrelated nouns and judged them independently. This study sets important boundary conditions for the limits of parallel processing of two words, but future work should attempt to generalize our model to conditions more similar to natural reading.

### Hemifield and hemisphere asymmetries

Another striking aspect of our data is that observers are much better at categorizing words to the right than left of fixation (Mishkin & Forgays, 1952; Nicholls & Wood, 1998; White et al., 2018). One potential explanation is the necessity of word-selective regions in the left hemisphere, which respond more strongly to words in the right than left hemifield (Fig. 2B). VWFA-1 in the right hemisphere may help recognize letter strings in the left hemifield (Le et al., 2017; Rauschecker et al., 2012; Strother et al., 2016). However, we found that the left hemisphere has three advantages: (1) there were more roughly three times as many voxels in left than right VWFA-1; (2) left VWFA-1 has two parallel channels, one for each hemifield; (3) only 1/3 of participants had a right VWFA-2 but all had a left VWFA-2, and the latter may contain the single channel through which all words must pass on the way to left-hemisphere language regions. Given that the word on the right of fixation has a stronger (and faster (Rauschecker et al., 2012)) bottom-up signal, it may automatically win a competitive normalization in left VWFA-2, blocking the left word at the bottleneck. Such a pattern has been observed with electrophysiological measurements in macaque face-selective brain regions: the contralateral face in a simultaneous pair dominates the neural response (Bao & Tsao, 2018). If left VWFA-2 behaves similarly, it could explain why accuracy for the left word is barely above chance in the distributed condition. In the focal cue condition, voluntary attention can shift the bias in that competition in favor of the left word, but only partially.

### Conclusion

The experiment reported here advances our understanding of the brain’s reading circuitry by mapping out the limits of spatially parallel processing and attentional selection. Surprisingly, parallel processing of multiple words extends from bilaterial retinotopic cortex into the posterior word-selective region (VWFA-1) in the left hemisphere. We propose that signals from the two hemifields then converge at a bottleneck such that only one word is represented in the more anterior VWFA-2. An important question for future work is whether similar circuitry applies to other image categories. Faces and scenes, for instance, are also processed by multiple category-selective regions arranged along the posterior-to-anterior axis in VOTC (Grill-Spector & Weiner, 2014). Recognition for each category might rely on similar computational principles to funnel signals from across the retina into a bottleneck, or written words might be unique due to their connection to spoken language.

## Materials and Methods

### Participants

15 volunteers from the university community (8 female) participated in exchange for a fixed payment ($20/hour for behavioral training and $30/hour for MRI scanning). All subjects gave written and informed consent in accord with the Institutional Review Board at the University of Washington, in adherence to the Declaration of Helsinki. All subjects were right-handed, had normal or corrected-to-normal visual acuity, and learned English as their first language. All scored above the norm of 100 (Mean ± SEM: 116.6 ± 2.7) on the composite Test of Word Reading Efficiency (TOWRE) (Torgesen, Rashotte, & Wagner, 1999).

The sample size was chosen in advance of data collection on the basis of a power analysis of a previous fMRI study (White et al., 2017). That study reported a mean selective attention effect of 0.1% signal change in retinotopic cortex. We simulated resampling those data to determine the number of participants required to detect an effect half as large with the same degree of noise. Thirteen subjects was the minimum required to reach 80% power. We rounded that number up to 15. Three participants had to be excluded and replaced. Two were excluded before any fMRI data collection because they broke fixation on at least 5% of trials during the behavioral training sessions. Another was excluded after one MRI session because he was unable to finish all the scans and failed to respond on 9% of trials (compared to the mean of 0.7% across all included subjects).

### Stimuli and task

During behavioral training, we presented stimuli with an Apple Mac Mini and a linearized CRT monitor with a 120 Hz refresh rate (1024 x 640 pixels). During MRI scanning, stimuli were generated with an Apple Macbook Pro and back-projected onto a fiberglass screen with a luminance linearized Eiki LCXL100 projector (60Hz; 1280 x 1024 pixels). The display background was set to the maximum luminance (90 cd/m^2^ during training and 2350 cd/m^2^ in the scanner). A fixation mark was present at the screen center throughout each scan: a black cross, 0.43 x 0.43 degrees of visual angle (°), with a white dot (0.1° diameter) at its center.

Word stimuli were drawn from a set of 246 nouns (see **Supplementary Tables S1** and **S2**). The nouns were evenly split between two semantic categories: “living” and “non-living”. The words were 4, 5 or 6 letters long, in roughly equal proportions for both categories. The mean lexical frequencies in the living and non-living categories were 18.6 and 14.3 per million, respectively. On each trial, one word was selected for each side, with an independent 50% chance that each came from the “living” category. The same word could not be present on both sides simultaneously, and no word could be presented on two successive trials.

Masks were strings of 6 random constants. Words and masks were presented in Courier font. The font size was set to 26 pt during training and 50 pt in the scanner, so that the size in degrees of visual angle was constant. The word heights ranged from 0.54° to 0.96° (mean = 0.78°), and their lengths varied from 2.5° to 4.2° (mean = 3.25°). All characters were dark gray on the white background (Weber contrast = −0.85).

Each trial **(Figure 1A)** began with a *pre-cue* for 1000 ms. The pre-cue consisted of two horizontal line segments, each with one end at the center of the fixation mark and the other end 0.24° to the left or right. In the *distributed cue* condition, both lines were the same color, blue or green. In the *focal cue* condition, one line was blue and the other green. Each participant was assigned to either green or blue and always attended to the side indicted by that color. After a 50 ms inter-stimulus-interval (ISI) containing only the fixation mark, the pre-masks appeared for 50 ms. The masks were centered at 2.75° to the left and right of fixation, and were followed by an ISI containing only the fixation mark (duration variable across subjects; see below). Then the two words appeared for 50 ms, centered at the same locations as the masks. The words were followed by a second ISI with the same duration as the first, and then post-masks (different consonant strings) appeared for another 50ms. After a third ISI, the *post-cue* appeared. This consisted of a green and a blue line, which in the single-task condition matched the pre-cue exactly. The line in the subject’s assigned color indicated the side to be judged. The post-cue remained visible for 1500 ms. During that interval the subject could respond by pressing a button (task description below). A 450 ms *feedback* interval followed the post-cue: the central dot on the fixation mark durned green if the subject’s response was correct, red if it was incorrect, and black if no response was recorded. Finally, there was a 650 ms inter-trial interval with only the fixation mark visible. Each trial lasted a total of 4 seconds.

The two ISIs between the words and masks had the same duration. That duration was adjusted for each observer during training to yield ∼80% correct in the focal-cue conditions, and then held constant in all conditions (mean = 84 ms). The duration of the third ISI, between the post-masks and the post-cue, was set such that the sum of all three ISIs was 200 ms.

The subject’s task was to report whether the word on the side indicated by the post-cue was a non-living thing or a living thing. The subject used their left hand to respond to left words and their right hand to respond to right words. For each hand, there were two buttons, the left of which indicated “non-living” while the right indicated “living.” In behavioral training, these four keys were “z” and “x” and “<” and “>”. In the scanner, the subject held a small button-box in each hand, each with two buttons on it.

In the focal cue condition, the pre-cue indicated with 100% validity the side to be judged on that trial. In the distributed cue condition, the pre-cue was uninformative, so the observer had to divide attention and try to recognize both words.

### Procedure

Each subject completed 3 or 4 one-hour training sessions before scanning. These began with the TOWRE test and task instructions, and the subject read the full list of words. Then they practiced the task, with the word-mask ISIs initially set well above threshold. The ISI was gradually shortened until accuracy in the focal cue condition settled at roughly 80% correct (averaged over left and right sides). Then the participant completed at least 14 “runs” (each run containing 21 trials of each condition). The ISI was the same in focal and distributed cue conditions, and was adjusted from day to day as necessary to maintain ∼80% correct in the focal cue condition.

Each participant then completed 3 MRI sessions. The first was for retinotopic mapping (see **Supplementary Methods**). In each of the 2^nd^ and 3^rd^ sessions, the participant completed 3 localizer scans (L) and 5 main experimental scans (M), in a fixed order: L, M, M, M, L, M, M, L.

### MRI data acquisition

Using a Philips Ingenia 3T scanner, we acquired anatomical images with a standard T1-weighted gradient echo pulse sequence (1-mm resolution). We acquired functional images with an echo planar sequence, with a 32-channel high-resolution head coil, a repetition time of 2 s, and an echo time of 25 ms. Thirty-five axial slices (80 × 80 matrix, 240 × 240 × 105-mm field of view, 0 gap) were collected per volume (voxel size: 3 × 3 × 3 mm).

### main experimental scans

Each six-minute scan contained 9 blocks of 7 trials. All trials in each block were of the same pre-cue condition: distributed, focal left, or focal right. During MRI scanning, there were 12-second blanks after each block, during which the participant maintained fixation on the cross. During behavioral training sessions, those blanks were shortened to 4 s. During the last two seconds of each blank, the pre-cue for the upcoming trial was displayed with thicker lines, to alert the participant.

### Localizer scans

We used localizer scans to define regions of interest (ROIs), presenting two types of stimuli one at a time at the same locations as the words in the main experiment **(Figure 1C)**. Each 3.4 minute localizer scan consisted of 48 four-second blocks, plus 4 s of blank at the beginning and 8 s of blank at the end. Every third block was a blank, with only the fixation mark present. In each of the remaining blocks, a rapid sequence of eight stimuli was flashed at 2 Hz (400 ms on, 100 ms off). Each block contained one of two types of stimuli: words or phase-scrambled word images, all either to the left or right of fixation (center eccentricity 2.75 deg). Therefore there were 4 types of stimulus blocks. Each scan contained 8 of each in a random order.

The words were drawn from the same set as in the main experiment, in the same font and size, but with full contrast. We created phase-scrambled images by taking the Fourier transform of each word, replacing the phases with random values, and inverting the Fourier transform. Each image was matched in size, luminance, spatial frequency distribution, and RMS contrast to the original word.

During the localizer scans, the subject’s task was to fixate centrally and press a button any time the black cross briefly became brighter. Those luminance increments occurred at pseudorandom times: the intervals between them were drawn from an exponential distribution with mean 4.5 s, plus 3 s, and clipped at a maximum of 13 s. Hits were responses recorded within 1 s after luminance increment; false alarms were responses more than 1 s after the most recent increment. An adaptive staircase (Kaernbach, 1990) adjusted the magnitude of the luminance increments to keep the task mildly difficult (maximum reduced hit rate = 0.8).

### MRI data analysis

We performed all analyses in individual brains, averaging only the final parameter estimates extracted from each individual’s regions of interest (ROIs). Using brainVoyager™ software, we first pre-processed each functional scan with: trilinear slice time correction; motion correction to the first volume of the first scan (trilinear detection and sinc interpolation); phase-encoding distortion correction, based on one volume collected in the opposite direction at the start of each session; and high-pass temporal filtering (cutoff: two cycles/scan). Each functional scan was co-registered with a high-resolution anatomical scan collected in the same session, which was itself co-registered with the anatomical scan from the retinotopy session.

We processed functional scans (combining across sessions) with the glmDenoise package in Matlab (Kay, Rokem, Winawer, Dougherty, & Wandell, 2013). The glmDenoise algorithm fits a general linear model to the task blocks and includes noise regressors estimated from voxels that were uncorrelated with the experimental protocol.

**Table 1** lists the numbers of voxels within each region of interest (ROI). A representative subject’s brain is illustrated in **Figure 2A**. ROIs in retinotopic areas (V1-V4, VO and LO) were defined from the localizer scan data by contrasting responses to scrambled words on the left minus scrambled words on the right. We defined each ROI as the intersection of voxels within that retinotopic area and the voxels that responded more to the contralateral stimuli at a conservative threshold of *p*<10^−6^.

**Table 1:** Mean (and standard error) numbers of voxels in each ROI. “Hem.” = hemisphere. Were able to define each ROI in all 15 subjects except right VWFA-1 (14/15 subjects) and right VWFA-2 (5/15 subjects).

|  | Left Hem. | Right Hem. |
| --- | --- | --- |
| V1 | 68 (5) | 68 (7) |
| V2 | 82 (8) | 90 (10) |
| V3 | 130 (15) | 126 (14) |
| V4 | 79 (10) | 90 (9) |
| VO | 42 (7) | 47 (10) |
| LO | 61 (15) | 82 (16) |
| VWFA_1 | 56 (7) | 17 (4) |
| VWFA_2 | 41 (5) | 19 (5) |

The visual word form area ROIs (VWFAs) were defined by the contrast of words - scrambled words, regardless of side, with the false discovery rate q<0.01. Voxels in all retinotopic regions were excluded from the VWFAs. Consistent with the emerging view that the visual word recognition system contains two separate regions in VOTC (at least in the left hemisphere), we separately defined VWFA-1 and VWFA-2 for each participant and hemisphere. In both hemispheres, VWFA-1 was anterior to area V4, often lateral to area VO, near the posterior end of the occipito-temporal sulcus (OTS). VWFA-2 was a second patch anterior to VWFA-1. In the left hemisphere, VWFA-2 was always anterior and/or lateral to the anterior tip of the mid-fusiform sulcus. Left VWFA-2 was usually also in the OTS, although in 4 cases it appeared slightly more medial, encroaching on the lateral fusiform gyrus. In a few cases, VWFA-1 and VWFA-2 were contiguous with each other at the chosen statistical threshold for the words - scrambled contrast. However, raising the threshold always revealed separate peaks, and VWFA-1 and VWFA-2 were defined to be centered around those peaks.

In the right hemisphere, there were considerably fewer word-selective voxels that met our statistical threshold (**Table 1**). We found VWFA-1 in 14/15 subjects, and VWFA-2 in 5/15 subjects. Three of those VWFA-2s were medial of the OTS, on the fusiform gyrus. Previous studies have also reported less word selectivity in the right hemisphere than in the left, and constrained to a single region (Grill-Spector & Weiner, 2014; Strother et al., 2016).

We analyzed the main experiment with GLM regressors for each type of block: distributed cue, focal cue left, and focal cue right. Blocks with one or more fixation breaks were flagged with a separate regressor. An average of 5 ± 1.4% of blocks were excluded in this way.

### Spatial encoding model

For each VWFA, the two-channel model assumes that there is one “spatial channel” for each word. Each channel is composed of the set of neurons that respond to words at its preferred location. Left and right channel response strengths are denoted *CL* and *CR,* respectively. Each voxel *i’s* response *D-,* is a weighted sum of the two channel responses:

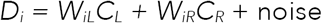

Weights *W*_*IL*_ and *W*_*IR*_ describe how strongly the two channels drive each voxel. We estimated those weights as the mean localizer scan responses to single words on the left (which evoke channel responses *C*_*L*_ = 1, *C*_*R*_ = 0) and single words on the right *(C*_*L*_ = 0, *C*_*R*_ = 1). That yielded a v-by-2 matrix *W* of voxel weights, where v is the number of voxels, and there is one column for each channel. Each condition of the main experiment (when two words are presented at once) produced a v-by-1vector *D* of voxel responses. The spatial encoding model can then be expressed as:

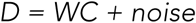

Linear regression gives the best-fitting estimate of C, a 2-by-1 vector of channel responses:

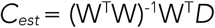

Note: (W^T^W)^−1^W^T^ is the Moore-Penrose pseudoinverse of the matrix W.

We compared that two-channel model to a one-channel model, in which each voxel *i* was assigned a single weight *W*_*i,avg*_ that was the average of its responses to left and right words in the localizer. Then there is a single channel that responds with strength *C*_*avg*_, such that:

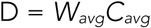

We then estimated *C*_*avg*_ using linear regression as well. For each model, in each subject and each ROI, we computed the proportion of variance explained, *R*^*2*^. We then adjusted each model’s *R*^*2*^for the number of free parameters, *p* (i.e. the number of channels):

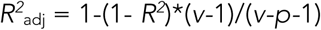

### Statistical analyses

We conducted linear mixed-effects models to assess the size and reliability of the effects of cue on BOLD responses (and estimated channel responses), and of how they differed across regions. In all models we included random intercepts across subjects, and when justified by a likelihood ratio test, we also included random slopes across subjects.

To assess the significance of the difference between pairs of conditions, we used bootstrapping: we built a distribution of 5000 means of N values resampled with replacement from the original sample of the N subjects’ differences. The two-tailed p-value is twice the proportion of bootstrapped means less than 0. At a significance cutoff of p=0.05, this approach is equivalent to regarding a difference as significant if the 95% confidence interval of differences excludes 0.

For more detail on the eye-tracking and retinotopy, see the **Supplementary Methods.**

## Acknowledgments

This work was funded by National Eye Institute grants K99 EY029366 to A.W., F32 EY026785 to A.W., and R01 EY12925 to G.B. and J.P. In addition, National Institute Of Child Health & Human Development grant R21 HD092771 to J.Y. and NSF/BSF BCS #1551330 to J.Y.

## Supplemental Information Appendix for

### Supplemental Methods

#### Eye-tracking and fixation control

We recorded the right eye’s gaze position with an Eyelink 1000 eye tracker (SR Research, Ottawa, Ontario, Canada). During behavioral training, we gave immediate feedback about fixation breaks. Each trial began only if the registered gaze position was within 0.75° horizontally and 3° vertically of the fixation mark. (We allowed more vertical tolerance to account for calibration drift and pupil size changes). We then averaged the gaze position over 10 samples to determine the initial fixation position. If, during the interval between the pre-cue offset and the post-mask offset, the estimated gaze position moved more than 1° horizontally or 2° vertically from the initial fixation position, the trial was immediately aborted. Text on the screen informed the participant that they had broken fixation and required a key-press to continue to the next trial. During MRI scanning, there was no such feedback about fixation breaks and trials continued at a constant pace. For one participant, technical errors prevented the recording of eye-tracker data during scanning, but their fixation control during training was excellent and they believed that their gaze position was still being monitored during scanning. For two other participants, eye-tracking failed on 1 and 3 scans, respectively (out of 10).

We detected fixation breaks in the recorded eye traces offline. For each scan, we defined the “central gaze position” as the median of all trials’ median gaze positions, each computed during the window between pre-cue and post-cue onsets. We cut out periods with blinks, ± 50 ms. We defined a fixation break a deviation >0.8° horizontally or >2.0° vertically that lasted more than 50ms and occurred between pre-mask onset and post-cue onset. Participants were excluded if they had fixation breaks on 5% or more of trials (applied to 2 participants after behavioral training). In behavioral training and in scanning, fixation breaks were detected on 2% and 1% of trials, respectively. Trials with fixation breaks were excluded from analysis of behavioral performance, and entire blocks with 1 or more fixation breaks were excluded from the MRI analysis.

#### Retinotopy

Each subject participated in a retinotopic mapping session. In each of six 4.2-minute scans, we presented one of three periodic stimulus types: a contracting ring, a rotating wedge, or alternating vertical/horizontal bow ties. All stimuli were composed of sections of radial checkerboards counter-phase flickering at 8 Hz. During each 256-s scan, the stimulus made eight ‘‘cycles’’ (rings contracting from 11.8 to 0.48 radius; wedge rotating clockwise one full circle; bow ties presented vertically then horizontally). The subject fixated a central white dot and pressed a button any time the dot briefly darkened or the checkerboard briefly dimmed in contrast. Using standard methods (Engel, Glover, & Wandell, 1997), we analyzed rings and wedge scans to identify the phase of the stimulus cycle that each voxel preferred, providing eccentricity and polar angle maps, respectively. We located the voxels representing the horizontal and vertical meridians via a general linear model (GLM) contrast of responses to the horizontal and vertical bow-tie stimuli. Using these activity patterns, we drew borders between retinotopic regions on each inflated cortical hemisphere. With these borders, we defined sets of anatomical voxels belonging to each retinotopic region. In some subjects VO1 was not clearly separable from VO2, and LO1 was not always separable from VO2. Therefore, we merged each pair of sub-regions (when there were two) into LO and VO.

**Table S1:** Stimulus set, Non-living category

|  |  |  |  |  |
| --- | --- | --- | --- | --- |
| vest | belt | chalk | penny | marble |
| sofa | shoe | flint | plate | carbon |
| glue | bath | villa | sword | blouse |
| gown | snow | dryer | wheel | velvet |
| coal | roof | scarf | knife | lounge |
| robe | coat | torch | clock | chapel |
| fork | moon | shack | pants | candle |
| sock | mask | cloth | slacks | canyon |
| mill | pipe | jeans | staple | nickel |
| coin | flag | lodge | mosque | cellar |
| silk | fuel | shelf | blazer | shorts |
| shed | soap | ridge | faucet | fridge |
| boot | iron | stove | pantry | pencil |
| dime | hinge | spoon | blinds | pillow |
| tire | whisk | attic | crater | carpet |
| pump | stair | bench | heater | wallet |
| lamp | clogs | skirt | bronze | closet |
| whip | shawl | drill | cradle | garage |
| barn | latex | frame | shrine | dollar |
| cave | satin | towel | drapes | toilet |
| fort | linen | motel | canvas | mitten |
| tube | cloak | steel | cement | sandal |
| sink | slate | cabin | bunker | castle |
| salt | manor | cable | dagger |  |
| sand | stool | couch | saloon |  |

**Table S2:** Stimulus set, Living category

|  |  |  |  |  |
| --- | --- | --- | --- | --- |
| bear | moth | heron | squid | iguana |
| bird | mule | horse | stork | insect |
| bull | newt | hound | tiger | jackal |
| bush | orca | human | trout | jaguar |
| carp | pine | hyena | tulip | lizard |
| clam | rose | koala | woman | maggot |
| crab | seal | lemur | whale | minnow |
| crow | slug | lilac | zebra | monkey |
| deer | swan | llama | amoeba | orchid |
| dove | toad | maple | baboon | oyster |
| duck | tree | moose | badger | parrot |
| fern | tuna | mouse | beaver | pigeon |
| fish | vine | otter | beetle | possum |
| flea | wasp | panda | cactus | python |
| girl | wolf | plant | canary | rabbit |
| frog | worm | raven | clover | salmon |
| goat | algae | robin | cougar | spider |
| hare | bison | shark | coyote | spruce |
| hawk | camel | sheep | donkey | turkey |
| kelp | cobra | shrew | falcon | turtle |
| lamb | daisy | shrub | ferret | walrus |
| lily | eagle | skunk | fungus | weasel |
| lion | finch | sloth | gerbil | willow |
| mole | goose | snail | gopher |  |
| moss | grass | snake | hornet |  |

## Supplemental Results

**Figure S1.**
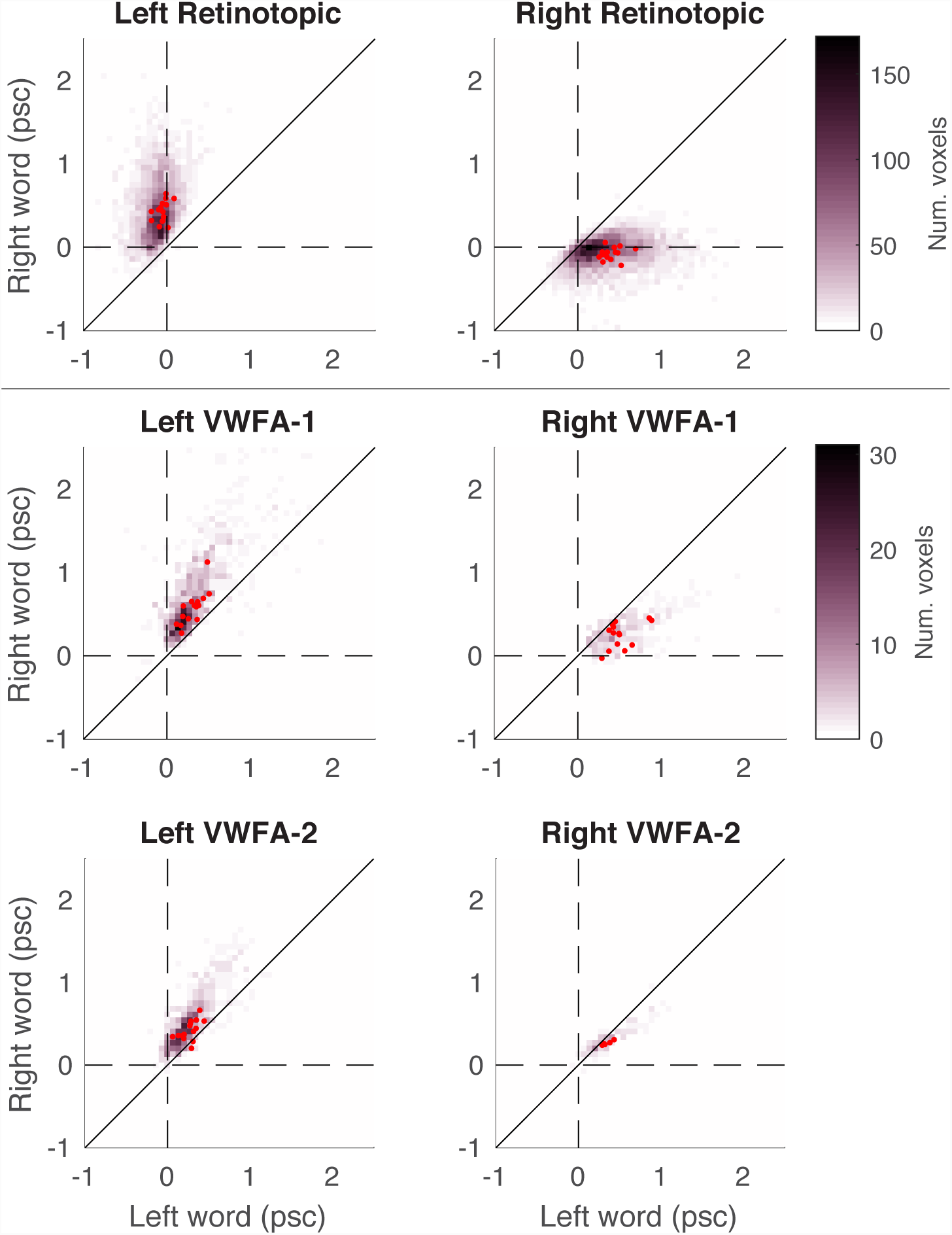
Voxel responses to words on the left and right of fixation in the localizer scans. The color of each point indicates the number of voxels with that combination of responses to words at the two locations. Red dots indicate the across-voxel mean for individual subjects. The top row contains the union of voxels from all retinotopic areas. The left column is for the left hemisphere, and right column for the right hemisphere.

**Figure S2.**
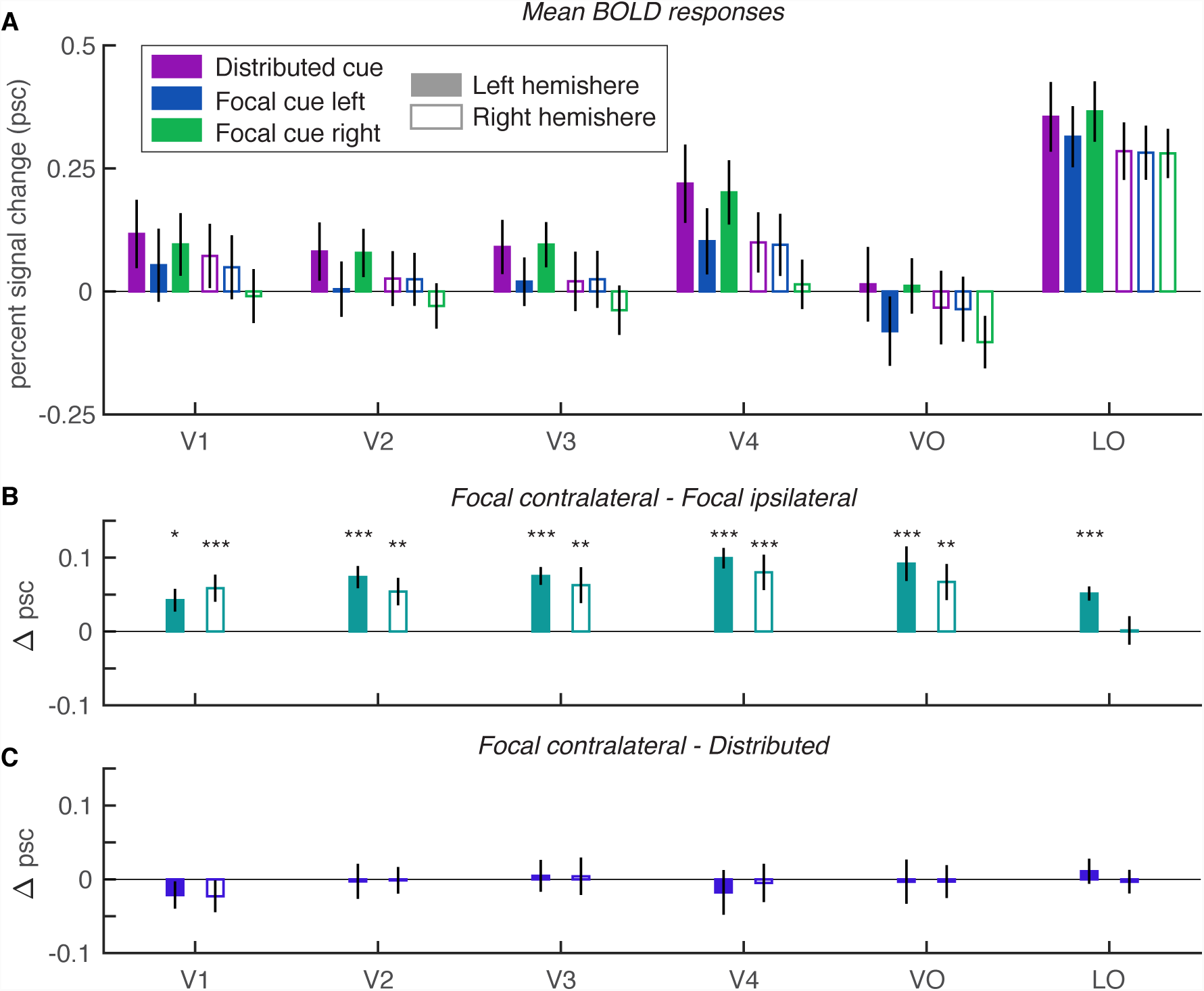
Mean BOLD responses and attention effects in retinotopic areas. **(A)** Mean BOLD responses and each ROI and hemisphere, divided by cue condition. **(B)** Mean selective attention effects: differences between responses when the contralateral vs. ipsilateral word was focally cued. **(C)** Mean divided attention effects: differences between responses when the contralateral word was focally cued vs. when both words were cued. All error bars = +/- 1 SEM. Asterisks indicate significant effects from bootstrapping: *** = p<0.001; ** = p<0.01; * = p<0.05.

**Figure S3.**
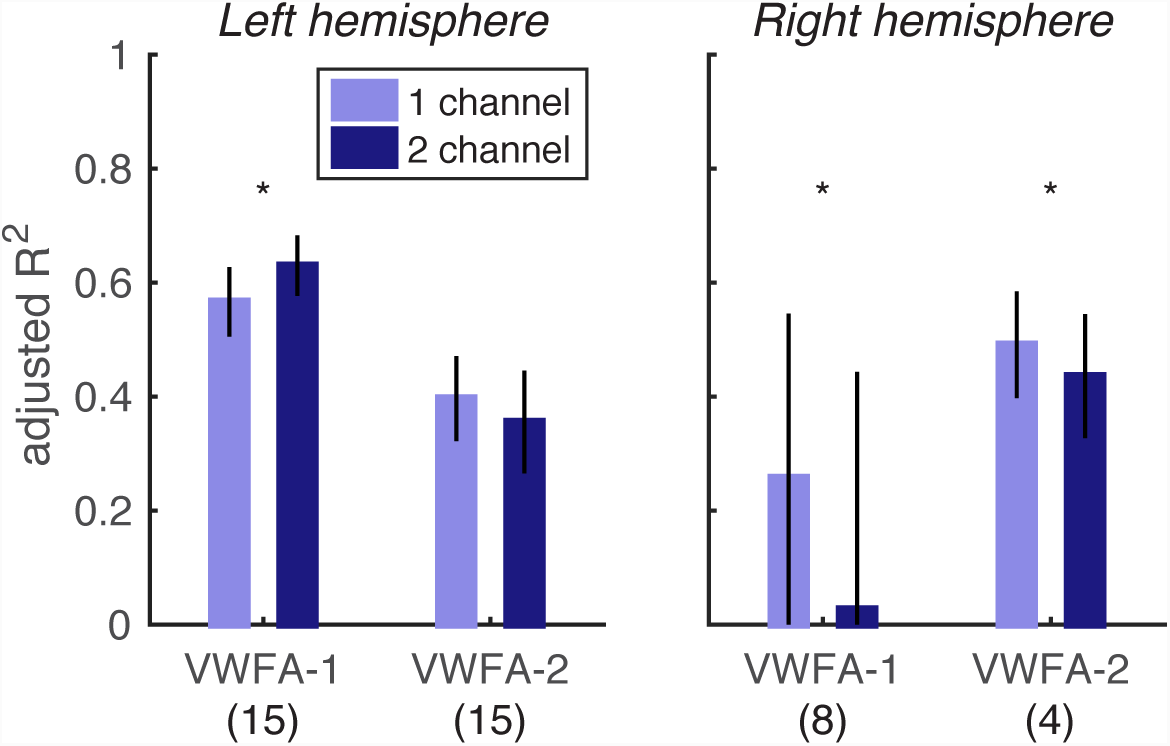
Adjusted r-squared values for the fit quality of the two-channel vs. the one-channel spatial encoding models. Asterisks indicate significant pairwise differences between the two models for each region (p<0.05 from bootstrapping). The numbers in parentheses at the bottom of the plot are the number of subjects that had enough voxels to compute adjusted R^2^ in each ROI.

**Figure S4.**
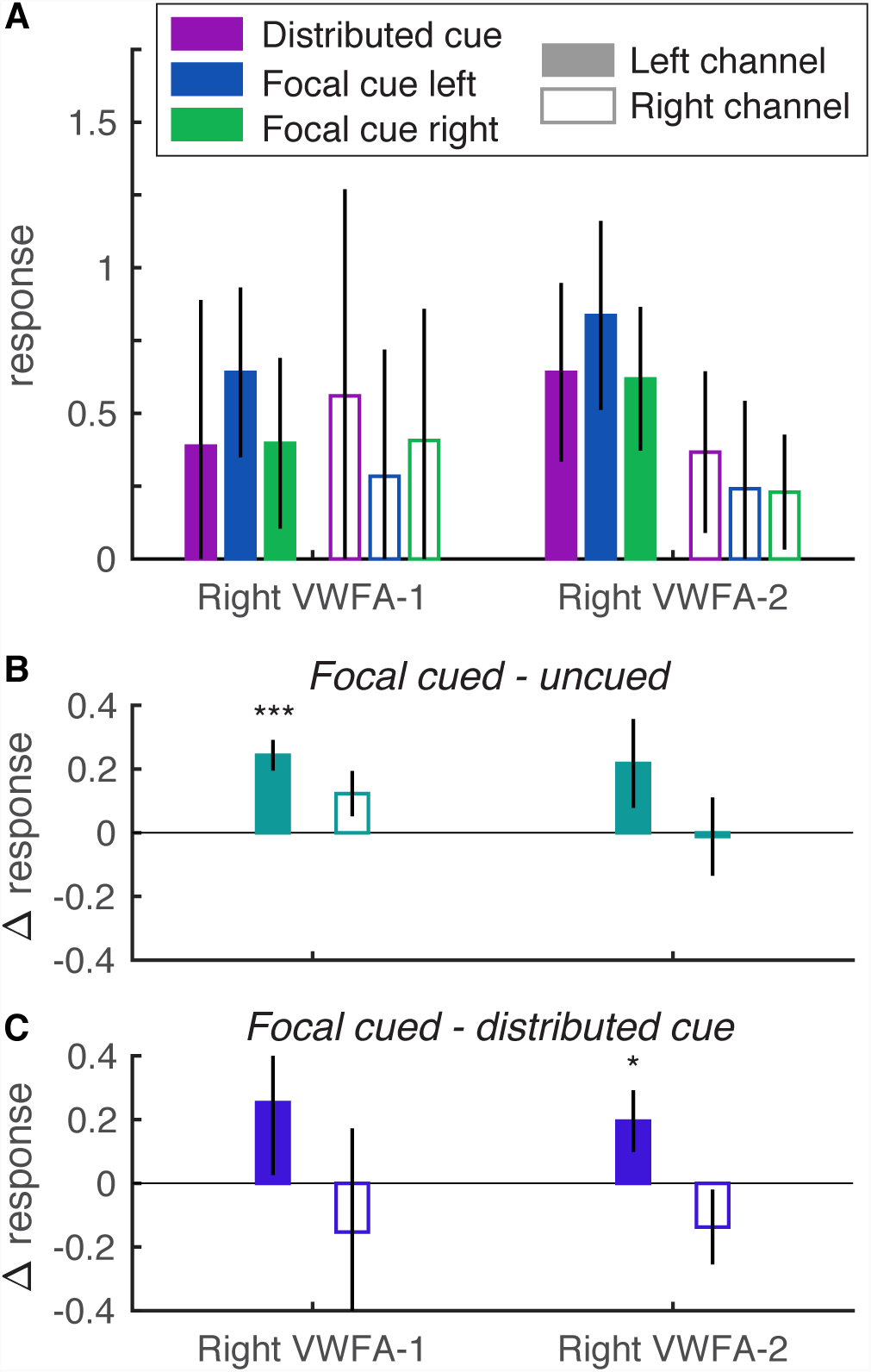
Estimated right hemisphere channel responses from the spatial encoding model. Error bars are ± 1 SEM. *** = p<0.001; ** = p<0.01; * = p<0.05. **(A)** Mean responses, separately for each ROI, channel, and cue condition. **(B)**Selective attention effects: the differences between each channel’s responses when its visual field location was focally cued vs. uncued. In Right VWFA-1 (14/15 subjects), there was a main effect of cue (mean = 0.18 ± 0.06; F(1,52)=12.0, p=0.001), no effect of channel (F(1,52)=0.06, p=0.80), but also an interaction (F(1,52)=5.94, p=0.018). The interaction indicates that the cue effect was bigger in the left channel (0.24 ± 0.05) than the right channel (0.12 ± 0.07), and only significant in the former. In Right VWFA-2 (5/15 subjects), there was no effect of channel (F(1,16)= 1.25, p=0.28), and no main effect of cue (F(1,16)=0.78, p=0.39), but an interaction (F(1,16)=44.8, p<10^−5^). The selective attention effect was bigger in the left channel (0.22 ± 0.14) than the right channel (−0.01 ± 0.12), but not significant in either. **(C)** Divided attention effects: the differences between each channel’s response when its location was focally cued vs. when both sides were cued. No effects of cue (focal cued vs. distributed) or channel (left vs. right) or interactions were significant, except for an interaction in right VWFA-2 (F(1,16)=5.55, p=0.031). The left channel was reduced by dividing attention (0.2 ± 0.10) but the right channel had the opposite effect (−0.14 ± 0.12). Note that 14/15 participants had a right VWFA-1 and only 5 had a right VWFA-2, and the one-channel model fit significantly better than the two-channel model in both regions.

**Figure S5.**
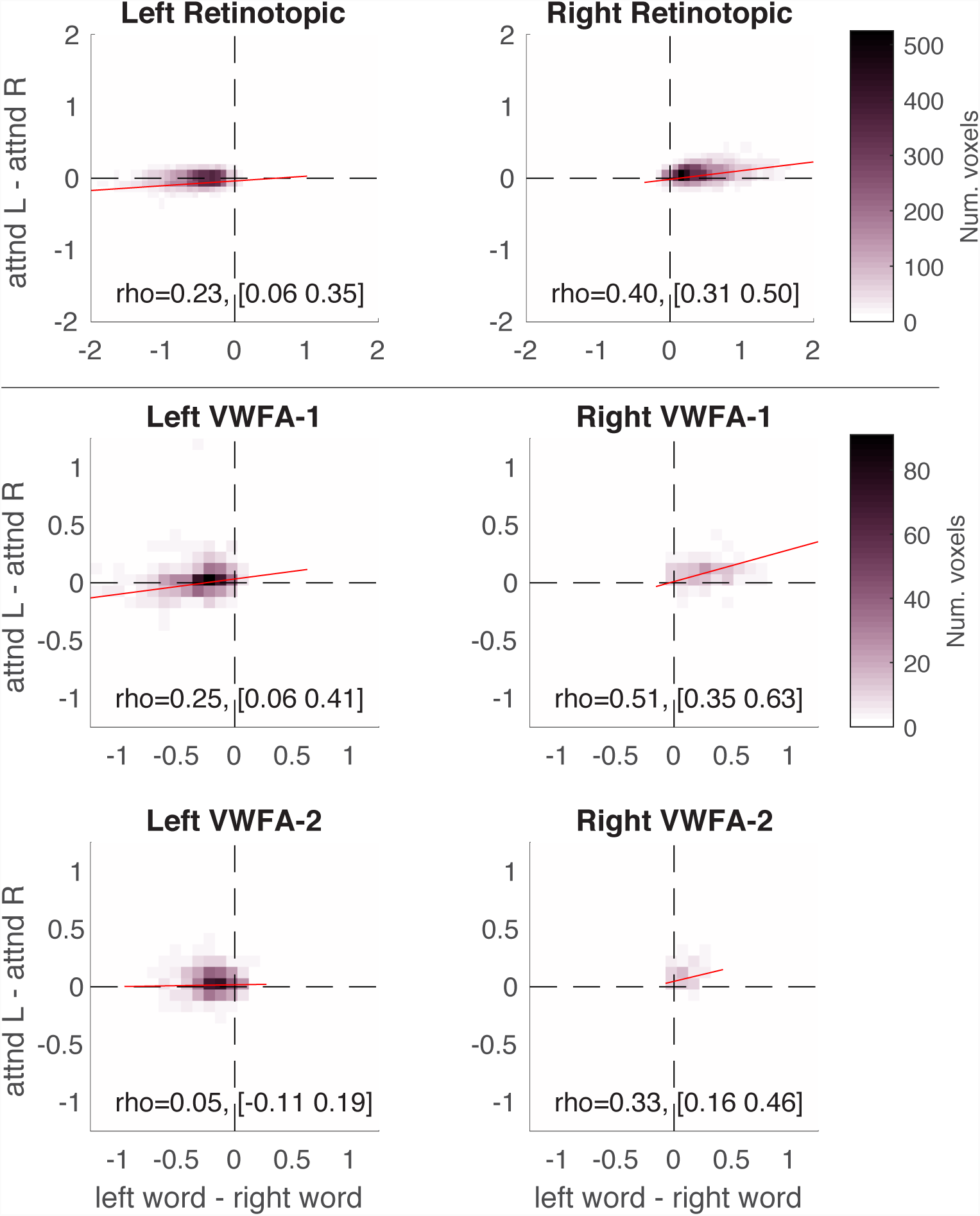
Voxel spatial vs. attentional selectivity. The x-axis is the difference in voxel responses between single words on the left and single words on the right in the localizer scans (spatial selectivity). The y-axis is the difference in voxel responses between the focal cue left and right conditions of the main experiment (selective attention effect). The color of each point indicates the number of voxels with each combination of those two differences. The text in each panel reports the across-subject mean correlation coefficient (rho) and it’s 95% confidence interval. The red line is the best-fitting linear mixed-effects model that included random effects of slope and intercept across subjects. The top row contains the union of voxels from all retinotopic areas. The left column is for the left hemisphere, and right column for the right hemisphere.

